# Resolving the developmental mechanisms of coagulation abnormalities characteristic of SARS-CoV2 based on single-cell transcriptome analysis

**DOI:** 10.1101/2023.03.13.532347

**Authors:** Xizi Luo, Nan Zhang, Yuntao Liu, Beibei Du, Xuan Wang, Tianxu Zhao, Bingqiang Liu, Shishun Zhao, Jiazhang Qiu, Guoqing Wang

**Author notes:** Correspondence to: Guoqing Wang, Jiazhang Qiu, Shishun Zhao. These authors contributed equally and are joint first authors on this work (Xizi Luo, Nan Zhang).

## Abstract

The COVID-19 outbreak caused by the SARS-CoV-2 virus has developed into a global health emergency. In addition to causing respiratory symptoms following SARS-CoV-2 infection, COVID-19-associated coagulopathy (CAC) is the main cause of death in patients with severe COVID-19. In this study, we performed single-cell sequencing analysis of the right ventricular free wall tissue from healthy donors, patients who died in the hypercoagulable phase of CAC, and patients in the fibrinolytic phase of CAC. Among these, we collected 61,187 cells, which were enriched in 24 immune cell subsets and 13 cardiac-resident cell subsets. We found that in response to SARS-CoV-2 infection, CD9^high^CCR2^high^monocyte-derived mø promoted hyperactivation of the immune system and initiated the extrinsic coagulation pathway by activating CXCR-GNB/G-PI3K-AKT. This sequence of events is the main process contributing the development of coagulation disorders subsequent to SARS-CoV-2 infection. In the characteristic coagulation disorder caused by SARS-CoV-2, excessive immune activation is accompanied by an increase in cellular iron content, which in turn promotes oxidative stress and intensifies intercellular competition. This induces cells to alter their metabolic environment, resulting in an increase in sugar uptake, such as that via the glycosaminoglycan synthesis pathway, in CAC coagulation disorders. In addition, high levels of reactive oxygen species generated in response elevated iron levels promote the activation of unsaturated fatty acid metabolic pathways in endothelial cell subgroups, including vascular endothelial cells. This in turn promotes the excessive production of the toxic peroxidation by-product malondialdehyde, which exacerbates both the damage caused to endothelial cells and coagulation disorders.

## Introduction

Severe acute respiratory syndrome coronavirus 2 (SARS-CoV-2) is a new human-infecting β coronavirus[1], and coronavirus disease 2019 has created an unprecedented global health crisis. Although most COVID-19 patients have relatively mild respiratory symptoms, headache, and fever[2], approximately 14% of patients will develop severe symptoms such as dyspnea, whereas 5% develop respiratory failure, shock, and multiple organ dysfunction, and may eventually die[3]. Severe COVID-19 is often accompanied by coagulation disorders, thromboembolism, and other complications[4], which are the main cause of multiple organ failure and death in patients with COVID-19[5]. Screening of case data from December 1^st^, 2019 to May 5^th^, 2021, revealed that the incidence of venous thrombosis in COVID-19 patients was as high as 20.9%, and that 85.9% of patients had at least one risk factor for venous thrombosis. Although the activation of coagulation is a hallmark of several infectious diseases, the pattern of coagulation activation in COVID-19 patients differs from that associated with sepsis[6, 7]. Indeed, sepsis-induced disseminated intravascular coagulation (DIC) is mainly manifested by a low platelet count, prolonged prothrombin time, and a reduction in antithrombin levels, whereas COVID-19 patients exhibit higher levels of fibrinogen and D-dimerization body levels, and there is comparatively little change in platelet counts, prothrombin time, or antithrombin levels. In addition, coagulation disorders induced by COVID-19 can occur in multiple organs, including the lungs, spleen, heart, and brain[8]. Despite anticoagulant prophylaxis, the probability of coagulopathy in patients with severe COVID-19 remains as high as 25% to 69%.

Accordingly to Conway et al., coagulation disorders in patients with severe COVID-19 will induce a cytokine storm, in which large numbers of cytokines stimulate circulating blood cells, causing blood cell activation and even apoptosis, thereby leading to thrombosis[9, 10]. In this regard, Bonaventura et al. have hypothesized that neutrophils are recruited for the release of neutrophil extracellular traps (NETs), which contain a range of prothrombotic molecules such as tissue factors, protein disulfide isomerase, factor XII, vWF, and fibrinogen. Extracellular DNA released by NETs promotes the direct activation of platelets, leading to thrombus formation[11]. Long et al. have proposed that monocytes produce the complement protein C5b-9, which enhances exposure of prothrombinase assembly sites on the surface of platelets by promoting the secretion of platelet factor V (FV) and assembly of a functional FXa/FVa complex, thus providing sites for thrombin propagation[12]. In previous studies, whereas researchers have focused on the relationships between certain cells and the development of coagulation disorders, comparatively little attention has been devoted to complex cellular composition and intercellular interactions.

In this study, we used single-cell sequencing technology to examine the cause of the coagulopathy characteristic of COVID-19 from the perspective of multiple cells and landscapes. On the basis of this single-cell sequencing, we analyzed data obtained for the right ventricular mononuclei derived from patients who died from CAC at different stages and from healthy donors and depicted the high-resolution transcriptome dynamic landscape and intercellular surface protein networks of each cell sub-population at different stages of CAC. We also investigated the correlation between coagulation mechanisms and immune activation, and constructed a polynomial calculation model, feature a scoring system, and a pathway activity algorithm to study the relationships among cellular iron levels, changes in oxidative stress, and metabolic reprogramming on cell competition and the exacerbation of coagulation disorders. On the basis of the data thus obtained, we provide a detailed characterization of the mechanisms associated with the development of coagulation disorders subsequent to SARS-CoV-2 infection, which will provide a valuable reference for the study of COVID-19-associated coagulopathy.

## Results

### Results 1: Comprehensive atlas of the coagulopathy characteristics of COVID-19

Our study included case information obtained for 170 severely ill patients with COVID-19. As shown in Supplementary Table 1, among these patients, the proportion over 66 years of age was 62.9%, and the proportions of patients with PT > 6% was 22.4%, prolonged APTT > 10 s was 40.3%, FIB > 1.9 g/L was 41.6%, and the ratio of D-dimer > 5.0 μg FEU/mL was 48.8%. These statistics indicate that most COVID-19 patients have coagulation abnormalities.

To systematically map a dynamic cellular atlas of CAC at different stages, for analysis, we used single-cell nuclear sequencing data obtained for the right ventricular free tissue of patients who died in the hypercoagulable phase of CAC, those in the fibrinolytic phase of CAC, and healthy donors. A corresponding flowchart is shown in Figure 1A. As shown in Figure1B, we performed an integrated quality control pipeline analysis of SnRNA-sequencing data of 17,694 nuclei obtained from healthy samples, 13,068 nuclei from those in the hypercoagulable phase of CAC, and 30,425 nuclei from those in the fibrinolytic phase. We generated gene-by-gene expression matrices, performed dimensionality reduction via Uniform Manifold Approximation and Projection (UMAP), graph-based clustering, and identified major cell subsets using eigengenes. We accordingly identified 10 major cell types, including immune cells, such as myeloid cells, NK cells, and subsets of cardiac constituent cells, including fibroblasts, cardiomyocytes, and adipocytes. Figure 1D shows the marker genes of each cell type used to cluster the cell sub-populations.

**Fig. 1.**
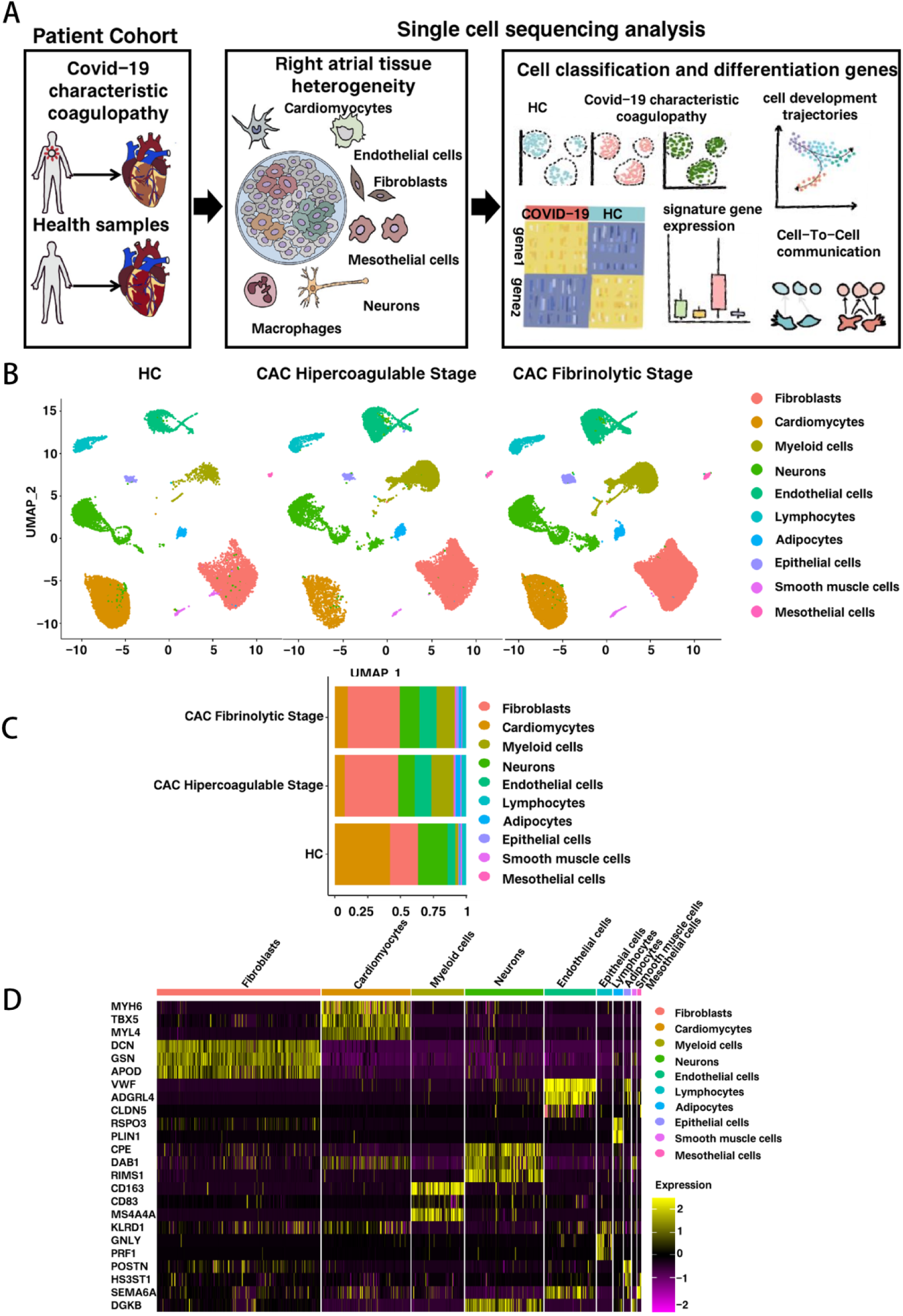
Analysis of the single-cell profiles in patients with coagulation disorders in COVID-19 and healthy controls. (A) Flow chart describing the overall experimental design of this study (B) Overview of cell clusters in the integrated single-cell transcriptome of 61,187 cells from patients with coagulopathy characteristic of COVID-19 and healthy controls (C) The proportion of cell subsets in CAC hypercoagulable phase, CAC fibrinolytic phase, and healthy controls (D) Heatmap of selected marker genes for cell subsets within cell lineages

We subsequently quantified the changes in cell-type ratio associated with the coagulopathy characteristics of COVID-19 patients. As shown in Figure 1C, in patients with coagulation disorders characteristic of COVID-19, the abundance of cardiac constituent cell subsets, such as cardiac cells and neuronal cells, was reduced, whereas the abundance of immune cells was increased, particularly in cases in which the myeloid cell subset was abnormally abundant. In addition, among the fibrinolytic phase samples, we detected a reduction in the proportion of immune cell subsets, such as those of myeloid and NK cells. On the basis of these data, we established that the immune microenvironment of patients with coagulation disorders characteristic of COVID-19 shows temporal differentiation.

### Results 2: Cardiac intercellular developmental transition in **the coagulation disorder characteristics of COVID-19**

In order to further study changes in the subpopulations of cardiac primary resident cells in COVID-19-associated coagulopathy, with the exception of except immune cells, we focused on changes in the subpopulations of cells in the cardiac cell atlas, combined with annotated genes, and further sub-clustered these, as shown in Figure 2A and 2C. Among these cell types, cardiomyocytes are mainly divided into two subgroups, ACTB^+^Cardiomyocytes and FHL2^+^Cardiomyocytes, the former of which are mainly involved in energy conversion and electrophysiological responses (Supplementary Figure 1H and 1J), whereas the latter are associated with retinoic acid and are involved in the differentiation and maturation of cardiomyocytes (Supplementary Figure 1I and 1K). As shown in Figure 2B, during the fibrinolytic phase of CAC, there is a reduction in the cell ratios of both these types of cardiomyocyte, which are more likely to initiate the Parthanatos mode of death. In addition, FHL2^+^Cardiomyocytes scored higher on death, indicating more severe damage (Figure 2E and 2F)[13]. During the fibrinolytic phase of CAC, the PI3K-AKT pathway is activated in both these cardiomyocyte subsets, which further promotes the formation of PIP1-TRAF2 death corpuscles and exacerbates cardiomyocyte death (Figure 2D)[14].

**Fig. 2.**
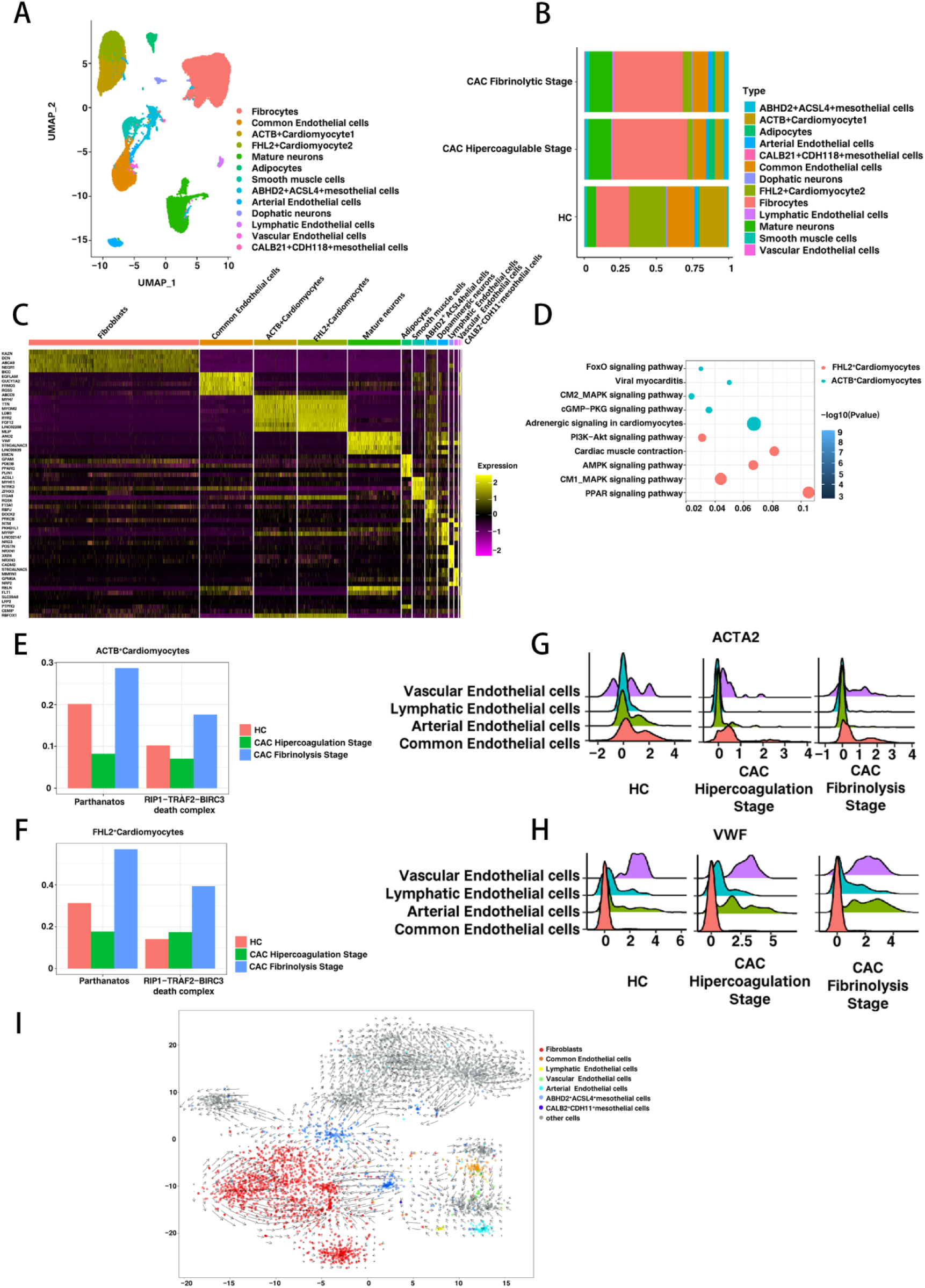
Characteristics of cardiac primary resident cell subsets in the characteristic coagulopathy of COVID-19. (A) UMAP projections highlighting clusters of primary resident cells in the heart. Clusters were named according to their cluster-specific gene expression patterns (B) Proportional distribution of cardiac primary resident cell subsets in the CAC hypercoagulable phase, CAC fibrinolytic phase, and healthy control hearts (C) Heatmap of selected marker genes for immune cell subsets within cell lineages (D) Differential pathway analysis of the CAC fibrinolytic phase in cardiomyocyte subsets (E-F) Death score assessment of cardiomyocyte subsets in the CAC hypercoagulable phase, CAC fibrinolytic phase, and healthy controls (G-H) Ridge plot of activated gene expression of endothelial cell subsets in the CAC hypercoagulable phase, CAC fibrinolytic phase, and healthy controls (I) Developmental trajectory analysis of cardiac primary resident cell subsets during the course of coagulopathy characteristic of COVID-19

In addition, we found that the proportion of fibroblasts involved in the coagulopathy characteristic of COVID-19 infection was enhanced compared with that in healthy controls, thereby providing evidence to indicate an increase in fibrosis during the course of COVID-19-associated coagulopathy. The developmental trajectory map shows that most of the ABHD2^+^ACSL4^+^mesothelial cells tend to develop toward fibroblasts during the coagulopathic process, which may account for the myocardial fibrosis in the COVID-19-infected group (Figure 2I). However, unlike ABHD2^+^ACSL4^+^mesothelial cells, we found that CALB2^+^CDH11^+^mesothelial cells do not develop into fibroblasts during the coagulopathic process. Interestingly, we found that whereas CALB2^+^CDH11^+^mesothelial cells were absent in healthy controls, they were enriched in both the hypercoagulable and fibrinolytic phases of CAC, which are the mainstays of the coagulopathy characteristic signature cell subsets following COVID-19 infection.

In the hypercoagulable phase of CAC, we detected the activation of endothelial cell subsets, predominately arterial and vascular endothelial cells, along with the high expression of a number of genes, including *VWF*, *ICAM1*, and *VCAM1*[15] (Figure 1H and 1G, Supplementary Figure 1E and 1G). Contrastingly, during the fibrinolytic phase of CAC, there was a reduction in the expression of endothelial-activated genes, such as *VWF* and *ICAM1*, in the arterial and vascular endothelial cells (Figure 1G, Supplementary Figure 1F), whereas the mesenchymal cell markers *ACTA2* and *AEB1* were highly expressed (Figure 1G, Supplementary Figure 1D). Similarly, we detected activation of the related pathways (NOTCH, TGF, and Wnt) involved in this process (Supplementary Figure 1A-1C). These observations thus provide evidence to indicate endothelial cells undergo transformation to mesenchymal cells, resulting in the EndoMT reaction[16, 17]. In the RNA rate graph shown in Figure 2I, a clear trend can be observed regarding the transformation of arterial and vascular endothelial cells.

### Results 3: During the course of CAC, there is a gradual **hyperactivation of the immune system**

To gain further insights into the immunological features of the coagulopathy characteristic of COVID-19, we performed sub-clustering of the immune cell lineages. According to the source and status of immune cell subsets and annotated genes, we identified a total of 24 immune cell subsets with specific transcriptional identities, as shown in Figure 3A and Supplementary Figure 2A. Among these, in healthy samples, the proportion of T cell subsets was 54.9% of the overall immune system, whereas in the coagulopathy characteristic of COVID-19, the proportion of T cell subsets was only 17% (Figure 3B). This type of reduction in the proportion of T cell subsets is a typical feature of changes in the immune microenvironment following COVID-19 infection[18].

**Fig. 3.**
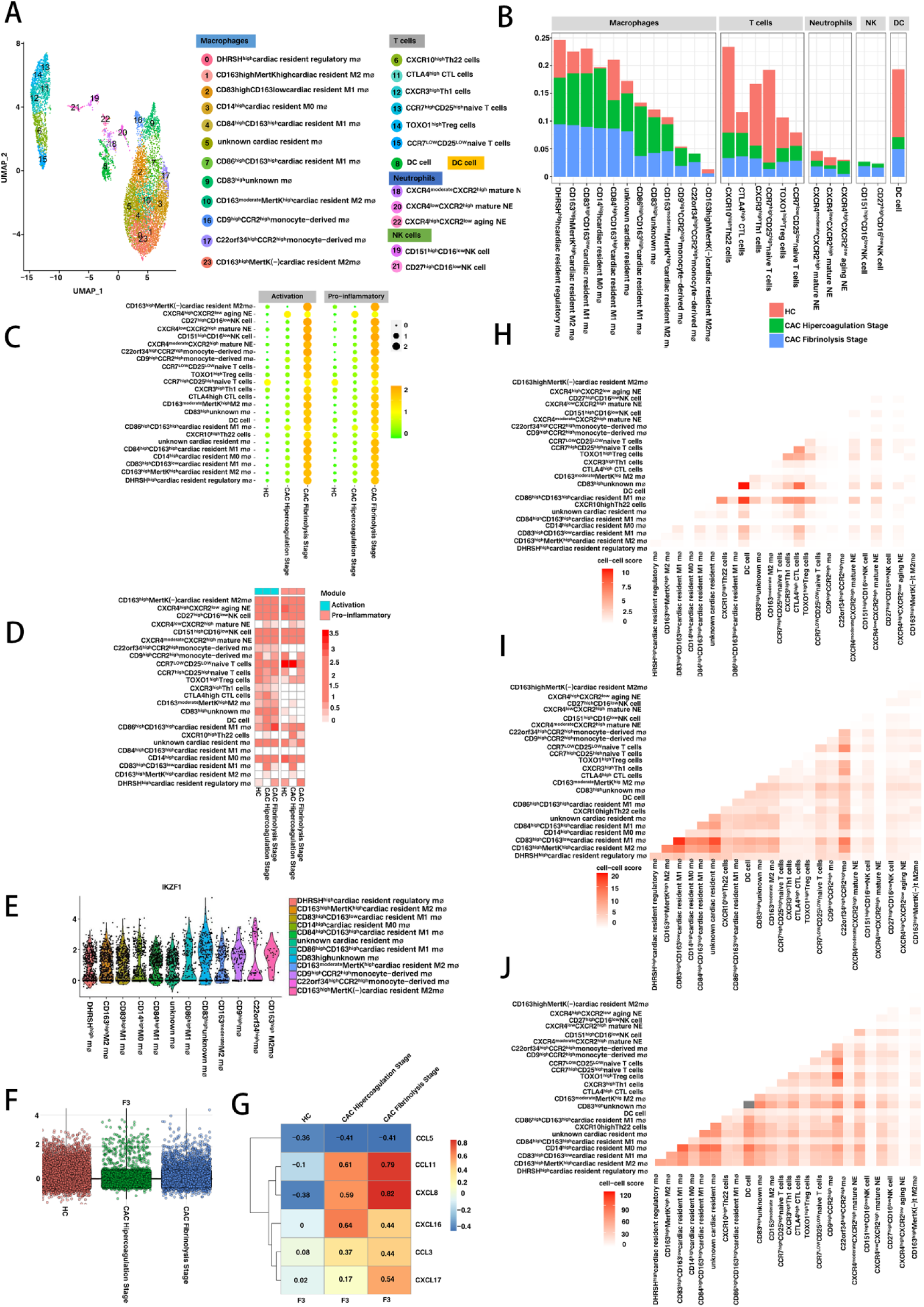
Immune signatures in the characteristic coagulopathy of COVID-19. (A) UMAP projection highlighting immune cell clusters. Clusters are named according to cluster-specific gene expression patterns (B) Proportional distribution of immune cell subsets in CAC hypercoagulable phase, CAC fibrinolytic phase, and healthy controls (C) Scores of immune cell subsets in immune activation and pro-inflammatory modules in CAC hypercoagulable phase, CAC fibrinolytic phase, and healthy controls (D) Evaluation of the correlation between immune cell subsets and immune activation and pro-inflammatory modules (E) Comparison of IKZF1 immune module expression in macrophage subsets during the fibrinolytic phase of CAC (F) Comparison of F3 expression in CAC hypercoagulable phase, CAC fibrinolytic phase, and healthy controls (G) Correlation analysis between F3 and selected chemokines (H-J) Analysis of the binding capacity between quantitative immune cell subsets in HC, CAC hypercoagulable phase, and CAC fibrinolytic phase

In the course of the coagulopathy characteristic of COVID-19, we found that compared with the hypercoagulable phase of CAC, the immune system was highly abnormally activated during the fibrinolytic phase. The scoring system of pro-inflammatory gene modules (including IL-1, TNF, IL-6, and IFN) and activation gene modules (including TYROBP, MYD88, IKBKB, and IFR5) increased in the fibrinolytic phase (Figure 3C). Moreover, it was found that the expression of chemokines continued to increase during the course of CAC coagulation disorder (Supplementary Figure 2B). During the fibrinolytic phase of CAC, CD14^high^cardiac resident M0 mø was significantly associated with proinflammatory modules, and CD9^high^CCR2^high^monocyte-derived mø was significantly associated with activation modules (Figure 3D). Among these, the CD14^high^cardiac resident M0 mø marked gene showed that the expression of MARCO (a scavenger receptor gene) is increased[19]. MARCO-mediated ligand delivery enhances intracellular TLR and NLR functions and activates TLR-induced DC cells. This provides further evidence to indicate that this is the main cause of immune dysregulation in the fibrinolytic phase of CAC. In normal tissues, levels of C22orf34^high^CCR2^high^monocyte-derived mø are barely detectable, although are slightly more common in the hypercoagulable phase of CAC, and substantially more abundant in the fibrinolytic phase. Therefore, this is considered a typical cell subset in the coagulation disorder process following COVID-19 infection. In the fibrinolytic phase of CAC, there are significant increases in the expression of the interferon gene TRF8, the chemokine pathway gene CXCL10, HCK, and other genes in C22orf34^high^CCR2^high^monocyte-derived mø, and the IKZF1 immune activation module score was higher than that of other macrophage subtypes (Figure 3E)[20]. These findings indicate that C22orf34^high^CCR2^high^monocyte-derived mø has an immune recruitment effect, and can be considered a hallmark cell subset of immune hyperactivation in the process of COVID-19-associated coagulopathy.

In order to further examine the dynamic characteristics and binding force changes among immune cells under conditions of CAC immune hyperactivation, we compared the expression profiles of immune cell subsets in HC and the hypercoagulable and fibrinolytic phases of CAC[21]. We also screened 82 pairs of surface proteins based on a proteomics database and literature review, and calculated the binding between immune cells according to the binding affinity of each protein pair and the expression of the genes encoding the pair of proteins (Supplementary Figure 3A). We also calculated the binding strength of each pair of cell surface proteins (Supplementary Figure 3B-3E). Among cardiac resident macrophages, expression of the MAPK activator gene MRAS and the MAPK pathway-related genes NR4A1 and IGF1 was significantly increased in CD163^high^MERK^high^M2 macrophages, which provides evidence to indicate that the MAPK pathway is activated. In monocyte-derived macrophages, we detected significant increases in peroxisome enhancer-activated receptor PPAR-related genes[22], PPARG, FABP4, and ALSL1 (Supplementary Figures 2E and 4B). These findings thus indicate that during the fibrinolytic phase of CAC, monocyte-derived macrophages modify their glucose and lipid metabolism based on PPAR, enhance glycolysis, and enhance macrophage polarization. During the fibrinolytic phase, there are increases in the interactions among DHRSH^high^cardiac resident regulatory mø, CD83^high^CD163^low^cardiac resident M1 mø, and CD84^high^CD163^high^cardiac resident M1 mø. In addition, we detected a significant enhancement of the interactions between CD9^high^CCR2^high^monocyte-derived mø and CXCR3^high^Th1 and CTLA4^high^CTL cells (Figure 3H-3J). Among the differential genes of CXCR10^high^Th22 cells, there was significantly up-regulated expression of the adhesion factors ITCA6 and PDGFD and the homing receptor CXCR6 (Supplementary Figure 2D), thereby indicating that during the fibrinolytic period of CAC, T lymphocytes are homing and activated. In contrast, there were reductions in the proportions of native T cells and down-regulated expression of the polarization-related genes PRKCQ and RBPT[23] (Supplementary Figure 2F), which tend to indicate a reduction in the proliferative capacity of T cells during the fibrinolytic phase. This is a common phenomenon observed in COVID-19 patients with coagulation disorders.

**Fig. 4.**
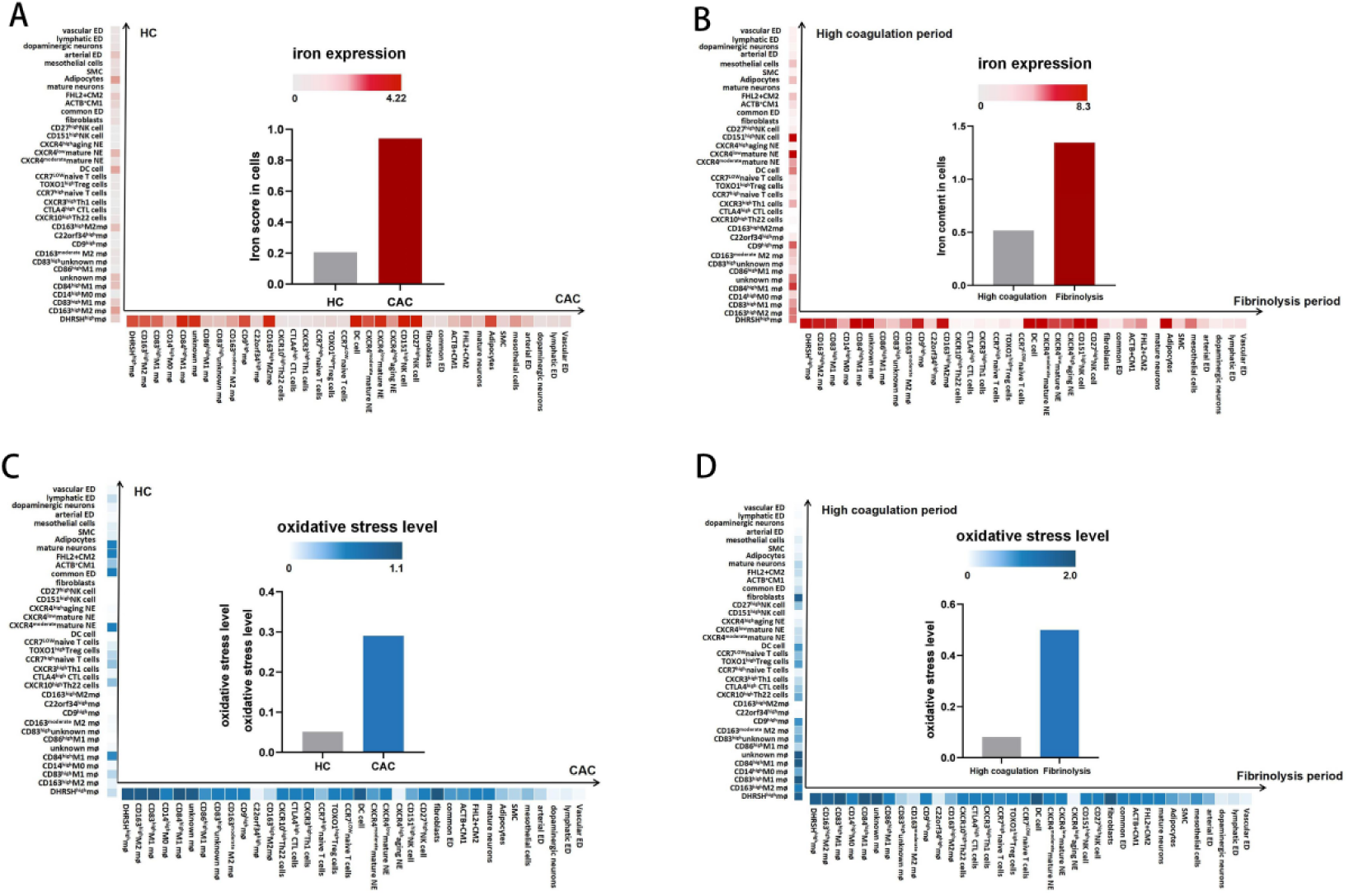
Elevated iron and oxidative stress levels following covid-19 infection. (A) Quantitative assessment of iron contents in HC and CAC samples (B) Quantitative assessment of iron contents during CAC hypercoagulable phase and CAC fibrinolytic phase (C) Quantitative assessment of oxidative stress in HC and CAC samples (D) Quantitative assessment of oxidative stress during CAC hypercoagulable phase and CAC fibrinolytic phase

Compared with healthy samples, we detected an enhancement in the correlation between chemokines such as CCL3 and CCL5 and tissue factor F3 in immune cells in samples with COVID-19-associated coagulopathy (Figure 3F and 3G). Compared with the hypercoagulable phase of CAC, there was a significantly higher correlation between chemokines and F3 in the fibrinolytic phase, and the expression of F3 was also significantly higher[24]. We found that during the process of CAC, the F3 gene in immune cells is completely expressed by CD9^high^CCR2^high^monocyte-derived mø cells during the fibrinolytic phase. The chemokine CXCR-GNB/G-PI3K-AKT signaling pathway is a key differential pathway in CD9^high^CCR2^high^monocyte-derived mø (Supplementary Figure 4A), and our analysis of the correlation between the CXCR-GNB/G-PI3K-AKT pathway and F3 at different stages of CAC in macrophages revealed a gradual increase in correlation (Supplementary Figure 2C). We accordingly speculate that in the immune thrombus induced by COVID-19 infection, CD9^high^CCR2^high^monocyte-derived mø initiate the extrinsic coagulation pathway by activating the CXCR-GNB/G-PI3K-AKT pathway.

### Result 4: Localized iron accumulation and oxidative stress **increase during the fibrinolytic phase of CAC**

In this study, we focused in particular on the changes in cellular iron content in COVID-19-associated coagulopathy As shown in Figure 4A and 4B, compared with healthy samples, patients swith COVID-19-associated coagulopathy had higher iron gene set scores, thereby indicating that hyperferritinemia occurs in these patients. Moreover, when focusing on the coagulopathy process, we found that iron contents in the fibrinolytic phase of CAC were significantly higher than those in the hypercoagulable phase.

The localized accumulation of iron will contribute to promoting the Fenton reaction, thereby resulting in an increase in ROS generation. To assess oxidative damage in the cellular microenvironment, we analyzed gene sets based on oxidative production (O), antioxidant capacity (R), oxidative stress level (OS). We quantified cellular oxidative stress levels by modeling polynomial functions, considering total oxidative capacity and antioxidant capacity, and assessing overall changes in oxidative stress. As shown in Figure 4C and 4D, compared with healthy samples, the COVID-19-associated coagulopathy samples were characterized by elevated levels of oxidative stress. Compared with hypercoagulable phase of CAC, higher levels of oxidative stress were detected in the fibrinolytic phase, thus indicating that during the development of coagulopathy characteristic of COVID-19, localized accumulation of iron in the CAC fibrinolytic phase triggers the Fenton reaction and exacerbates cellular oxidative damage.

### Result 5: Cell competition and cell injury are exacerbated by **elevated levels of iron and ROS**

In response to a combination of SARS-CoV-2 invasion and oxidative stress, cells undergo metabolic reprogramming to adapt to the modified tissue environment. During the development of coagulopathy characteristic of COVID-19, we detected an increase in cell microenvironment oxidative stress during the fibrinolytic phase of CAC, thereby promoting an increase in intercellular competition[25].

As shown in Figure 5A-5C, we observed an increase in the uptake of sugar by all immune cells during the fibrinolytic phase of CAC, and compared with the hypercoagulable phase, we detected an increase in the phosphatidylinositol metabolism of M1 macrophages during the fibrinolytic phase. This enhanced metabolism of phosphatidylinositol enables M1 cells to obtain larger amounts of phospholipids and contributes to the formation of phagosomes, thereby enhancing the phagocytosis of viruses. CXCR4^moderate^CXCR^2high^ mature NE and CXCR4^low^CXCR2^high^ mature NE increase fatty acid metabolism and the lysine degradation pathway, in the latter of which, the acetyl-CoA produced promotes the TCA cycle of neutrophils and provides energy for the cellular competition of neutrophils. In addition, during the fibrinolytic phase of CAC, DC cells are characterized by increases in the biosynthesis of pantothenate and CoA, which are key precursors for biosynthesis, the elevated levels of which can promote the TCA cycle and phospholipid synthesis. Nitrogen metabolism in NK cells is also elevated, and given the toxic nature of nitrogen metabolites, this can perturb the immune function of NK cells. In the immune microenvironment, wherein virus invasion pressure and a stressful environment coexist, immune cells act synergistically to resist the viral attack, and in response to oxidative stress, there is an increase in intercellular competition[26].

**Fig. 5.**
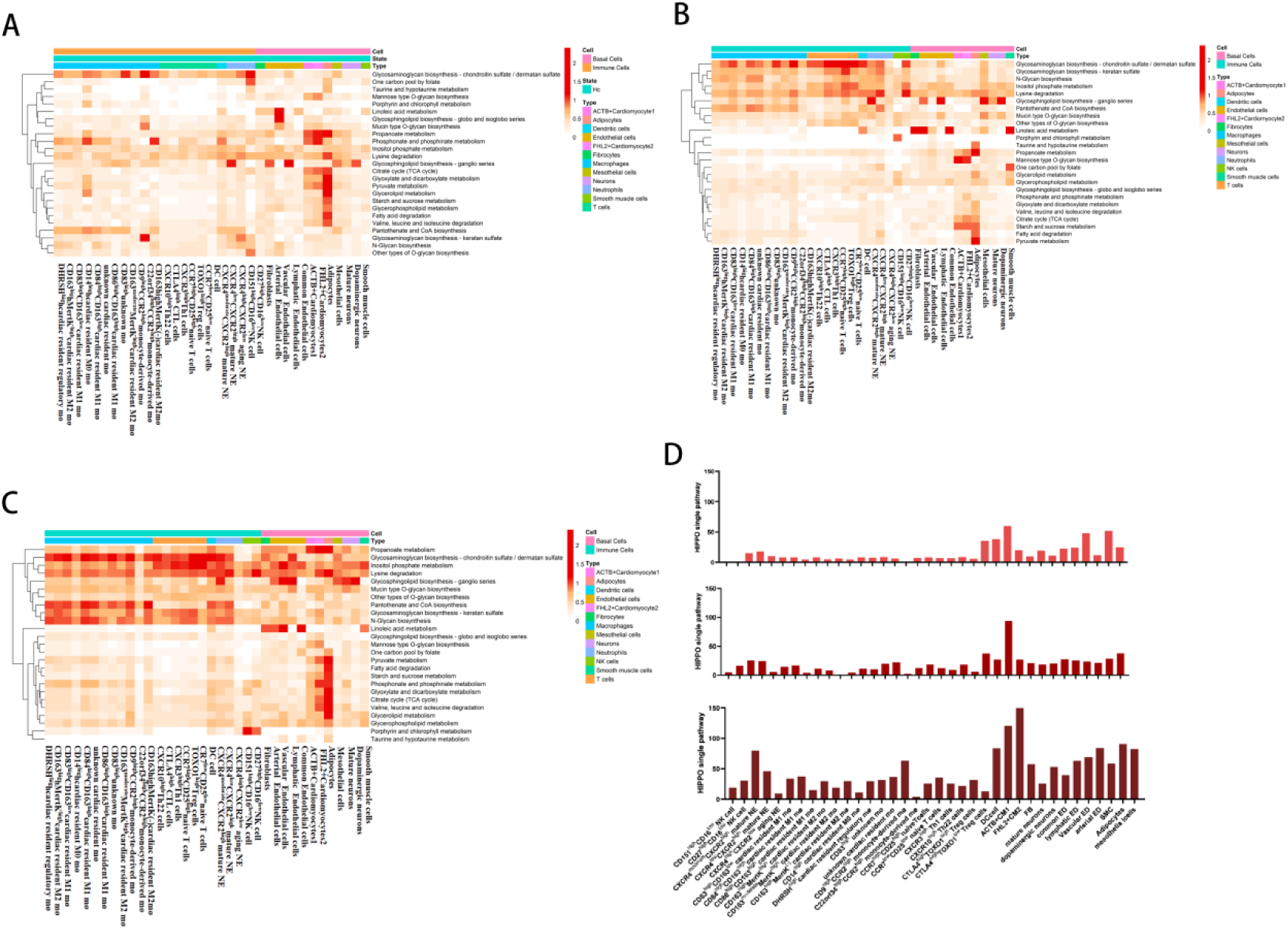
Cell competition and endothelial loss in coagulation abnormalities in COVID-19. (A-C) Analysis of metabolic reprogramming in HC, CAC hypercoagulable, and CAC fibrinolytic phases (D) Warts–Hippo pathway scores in the HC, CAC hypercoagulable phase, and CAC fibrinolytic phase

Stromal cells also initiate a unique competitive approach to dealing with oxidative stress. During the fibrinolytic phase of CAC, glycan biosynthesis is elevated in fibroblasts. Glycans are cell surface molecules that can respond to environmental stress by altering their glycosylation status without undergoing structural modification. In FHL2^+^CM, the TCA cycle is accelerated in adipocytes to meet cellular energy demands. In addition, the enhanced degradation of the amino acids valine, leucine, and isoleucine in adipocytes provides a source of TCA cycle metabolic mediators. As shown in Figure 5D, activation of the Warts-Hippo pathway, a cell-competitive marker pathway, was significantly increased during the fibrinolytic phase compared to the hypercoagulable phase of CAC[27]. These finding provide further evidence to indicate that intercellular competition is enhanced during the fibrinolytic phase of CAC.

During the fibrinolytic phase of CAC, endothelial cells, such as lymphatic and vascular endothelial cells, are characterized by increased levels of unsaturated fatty acid and lipid metabolism, resulting in the production of malondialdehyde. As shown in supplementary Figure 5A-D, we detected increase in the levels of TBXAS1, PTGS2, and PTGS1 in lymphatic and vascular endothelial cells and other cells during the coagulation disorder of CAC, and observed a corresponding elevation in the levels of malondialdehyde in endothelial cells[28],. We therefore speculate that elevated levels of iron and ROS contribute to increases in the levels of malondialdehyde in endothelial cells[29], which thereby exacerbates endothelial injury.

## Discussion

Coagulation disorders and thromboembolism are important causes of increased mortality in patients with severe COVID-19. We found that in response to SARS-CoV-2 infection, CD9^high^CCR2^high^monocyte-derived mø promotes immune hyperactivation and activates the PI3K-AKT pathway, which promote cytokine secretion and initiate the exogenous coagulation pathway, thereby leading to CAC. During the development of CAC, the immune system is gradually over-stimulated, and the binding force between immune cells is progressively enhanced. Among these cells, C22orf34^high^CCR2^high^monocyte-derived mø is are hallmark subset of cells associated with coagulopathy characteristic of COVID-19, and these cells play roles in immune recruitment. CD9^high^CCR2^high^monocyte-derived mø are a further important subset of cells that promote immune hyperactivation. In addition, in coagulation disorders characteristic of COVID-19, CD9^high^CCR2^high^monocyte-derived mø are characterized by the high expression of the exogenous coagulation initiation factor F3. Furthermore, among the differential pathways of CD9^high^CCR2^high^monocyte-derived mø, the pro-chemokine CXCR-GNB/G-PI3K-AKT signaling pathway shows a strong correlation with the procoagulant factor F3 in the coagulation disorders characteristic of COVID-19. Consequently, we believe that in the immune thrombus that develops following COVID-19 infection, activation of the extrinsic coagulation pathway by CD9^high^CCR2^high^monocyte-derived mø is the main process contributing to the development of coagulation disorders.

Excessive immune activation is typically associated with elevated iron levels, resulting in heightened levels of intercellular competition and greater endothelial cell damage, which are prominent factors contributing to coagulation abnormalities. We detected elevated levels of iron in cells during immune hyperactivation caused by COVID-19 infection. Consistently, the findings of studies conducted at the Athens National and Kapodistrian University School of Medicine in Greece[30] and Istanbul Medipol University[31] revealed the that patients with severe COVID-19 infection are often characterized by hyperferritinemia. We believe that elevated levels of cellular iron trigger the Fenton reaction, resulting in an increase in the production of ROS. By quantifying the level of oxidative stress, we also further confirmed the gradual increase of cellular oxidative stress in processes associated with coagulation disorders characteristic of COVID-19. In response to virus invasion and elevated levels of iron and ROS, cells modify their metabolic pathways in order to cope with an intensification of intercellular competition. During the fibrinolytic phase of CAC, there is an increase on the uptake of sugar by cells, along with an acceleration of the metabolism of linoleic acid and α-linolenic acid in endothelial cells. Higher levels of α-linolenic acid metabolism can in turn enhance cellular activity, up-regulate genes associated with lipid metabolism, and promote fatty acid oxidation. We also detected an enhancement of TCA cycle activity in adipocytes and FHL2^+^Cardiomyocytes. Elevated iron content not only has the effect of intensifying cellular competition but also exacerbates endothelial cell damage. During the fibrinolytic period of CAC, endothelial cells are characterized by an excessive activation of pathways associated with the metabolism of unsaturated fatty acids, thereby resulting in the excess generation of the peroxidation degradation product malondialdehyde in endothelial cells. Elevated levels of malondialdehyde will alter the permeability of endothelial cell membranes, thereby promoting cell double membrane rupture, and thus exacerbating endothelial injury. A number of previous studies have reported significantly higher levels of malondialdehyde in patients with venous thrombosis compared with those recorded in normal group subjects, and this metabolite has been established to play key roles in the development of thrombosis and coagulation disorders.

Our findings in this study provide evidence to indicate that cellular levels of ferritin are positively associated with the coagulopathy characteristic of COVID-19, that is, elevated iron levels exacerbate coagulation disorders in patients with COVID-19. Consequently, real-time detection of changes in iron concentrations in patients with COVID-19 and the maintenance of iron levels within the normal range could effectively contribute to controlling the occurrence of coagulopathy characteristic of COVID-19[32]. In conclusion, in this study, we analyzed the causes of COVID-19-associated coagulopathy, and characterized the dynamic changes of cells and ecological environment at different stages of this disorder. Our findings provide valuable insights that will contribute to the development of therapeutic approaches for the control and treatment of COVID-19-associated coagulation disease.

In this study, we analyzed the right ventricular single-cell nuclear sequencing data obtained for healthy donors and patients who had died at different stages of COVID-19-associated coagulopathy. We aimed to determine the causes of abnormal coagulation in COVID-19 patients and to provide detailed insights into the progression of coagulopathy in these patients.

## Methods

### Data collection

Raw data for the normal group were obtained from the Human Cell Atlas (HCA) Data Coordination Platform (DCP) (accession number: ERP123138; https://www.ebi.ac.uk/ena/browser/view/ERP123138). Sample data for COVID-19 patients with coagulation abnormalities were obtained from the Single Cell Portal (https://singlecell.broadinstitute.org/single_cell/study/SCP1464/single-nuclei-rna-sequencing-analysis-of-cardiac-microthrombi-in-fatal-covid-19) and Gene Expression Omnibus (https://www.ncbi.nlm.nih.gov/geo/) obtained GSE185457. For the normal group, we selected samples D2, H2, and H3, and among the samples for COVID-19 patients with coagulopathy, we selected GSM5615618, GSM5615619, GSM5615620, GSM5615621, GSM5615622, GSM5615623, and GSM5615624. Following data cleaning and sample merging, and the data were integrated into a dataset for analysis.

### Clinical characteristics after data pooling

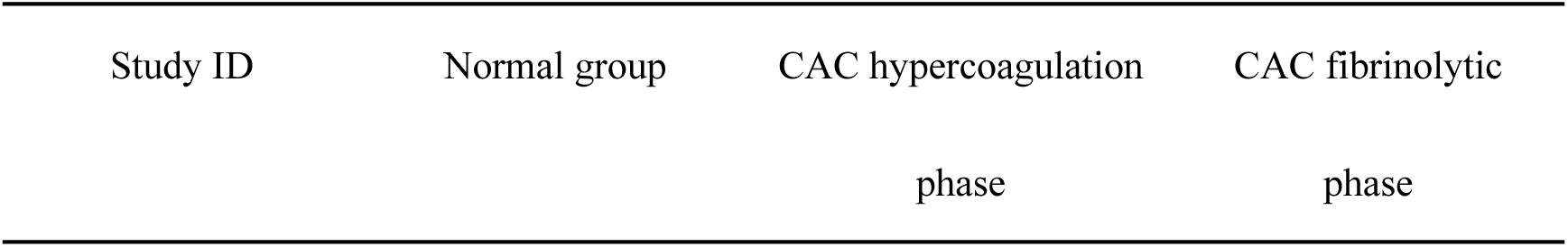
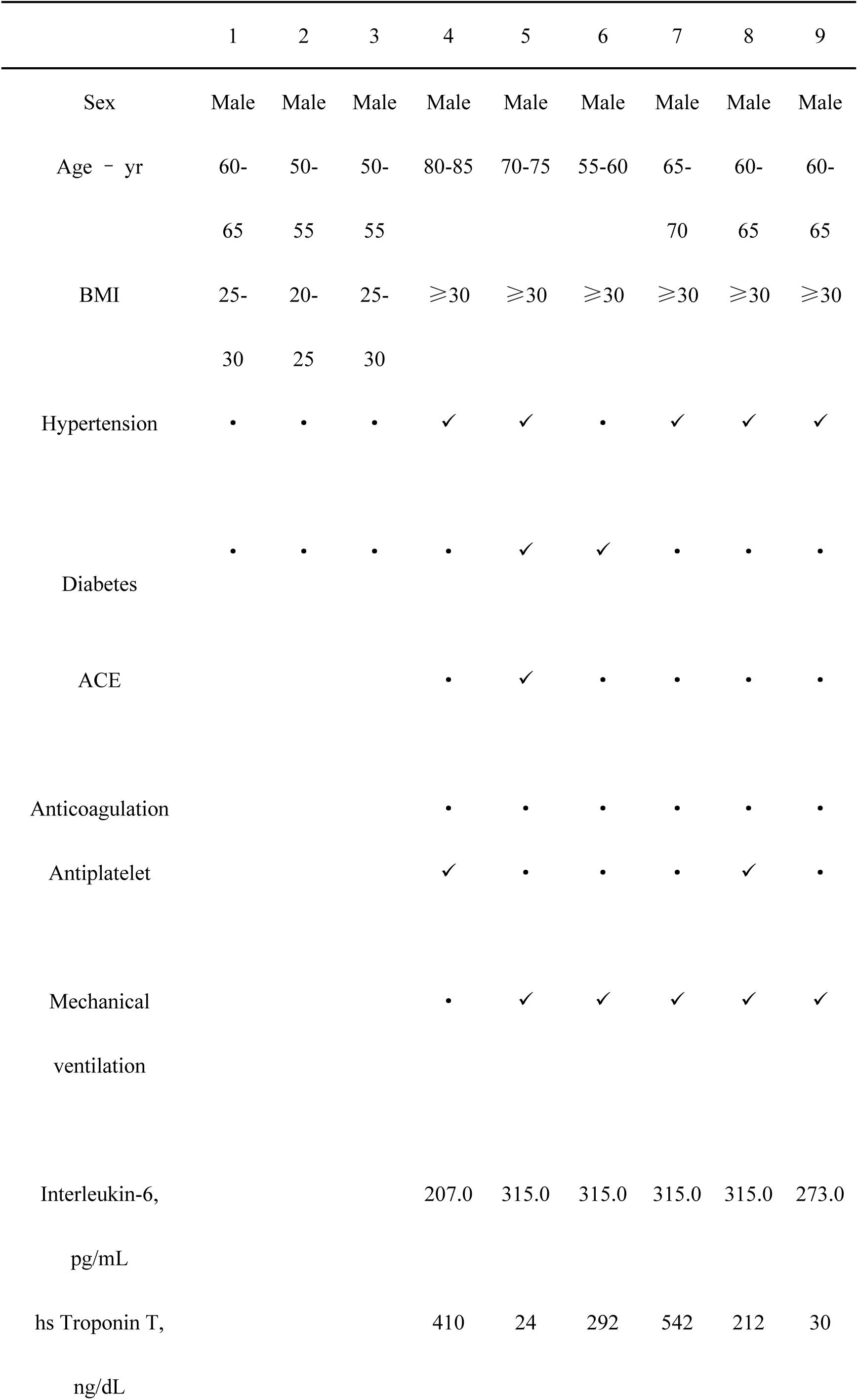
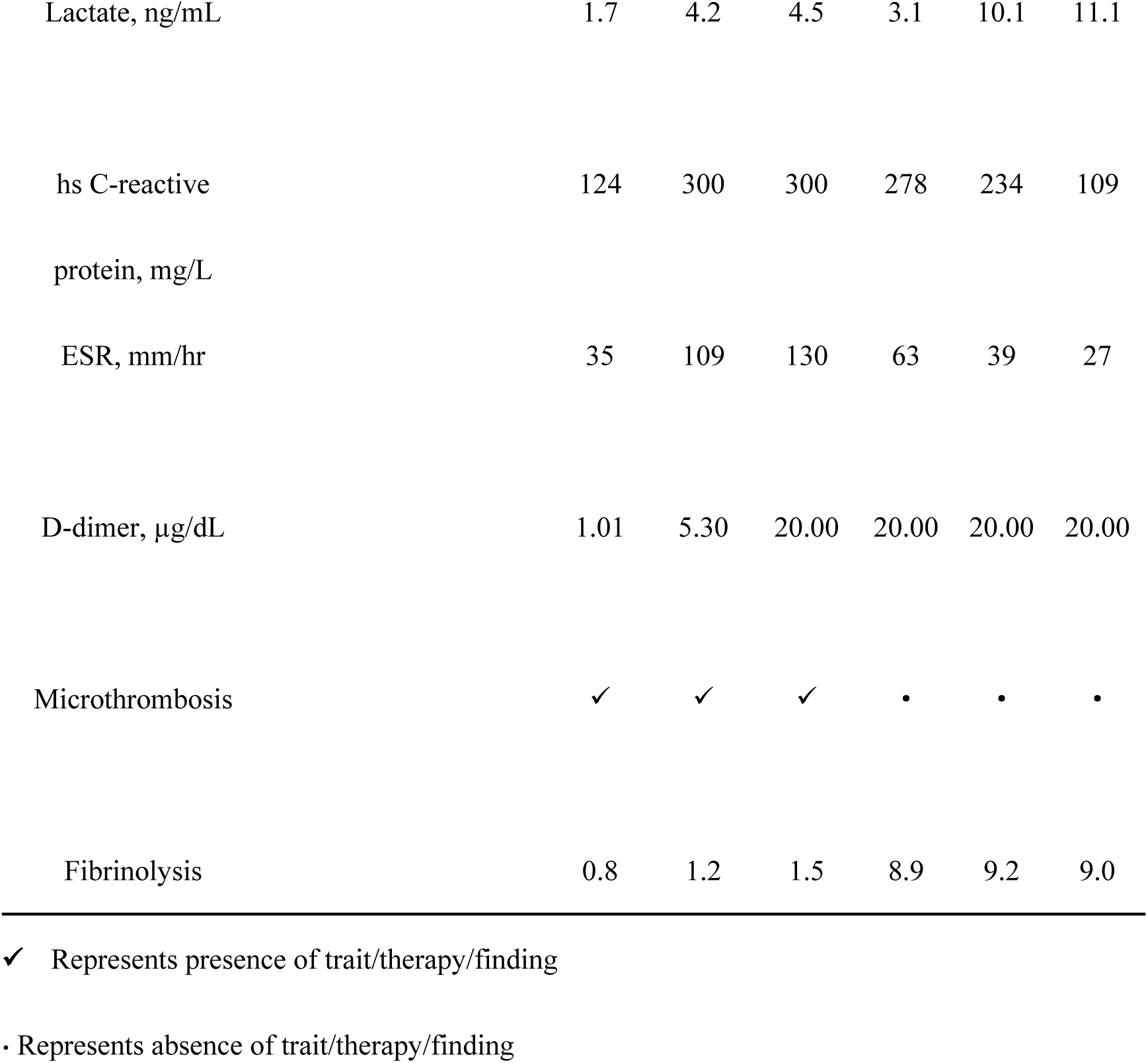

### Cell clustering

We used the SingleR package for machine annotation and the Cellmarker website and the latest relevant literature for further manual annotation. The first round of annotation identified 10 major cell types, including lymphocytes, myeloid cells, fibroblasts, cardiomyocytes, and endothelial cells. To identify clusters within each major cell type, we performed a second round of clustering on immune cells versus cardiac proto-resident cells, which followed the same process as the first round, starting with the low-level concordance output of highly variable genes selected as described above, with a resolution ranging from 0.3 to 1.8. We used the limma package for cell scale change calculations.

### Differential expression and gene ontology enrichment **analysis**

To investigate the effect of the presence of viral RNA on cells, we identified differentially expressed genes using the limma package. Functional enrichment was determined for each group of cells using the R package Profiler with default parameters, and the data were visually displayed using the ggplot2 and Heatmap packages.

### Inflammation and cytokine correlation scores

We downloaded an immune activation, immune recruitment, inflammation, cytokine-related gene sets from MSigDB, and scored related gene sets based on the total gene expression and cell number in the gene set.

### Immune cell surface protein interactions

We screened high-resolution proteomic datasets to extract cell surface protein pairs, and searched Google and PubMed for their names and applicable synonyms using standardized search terms, including <“protein name” and (binding or interaction or affinity) > and <“protein name” and (SPR or kinetics)>, to obtain the inter-protein pair affinity KD^3D^. A cell surface protein net was obtained by combining the protein pair affinity with the gene concentration encoding the protein pair using the following formulae:

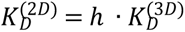

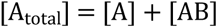

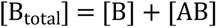

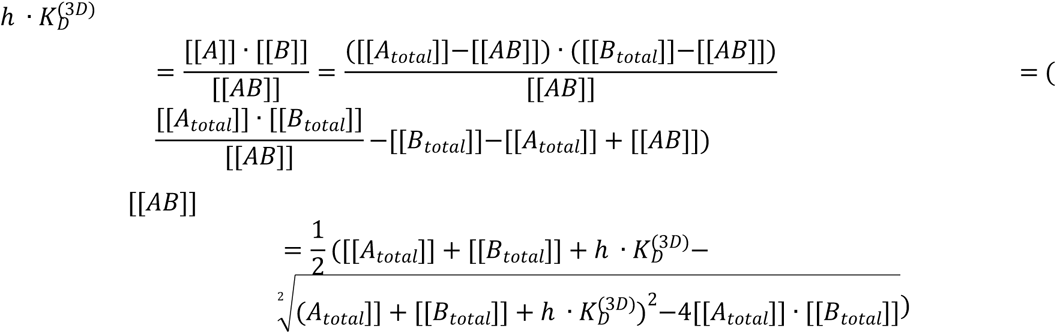

### Quantification of oxidative stress

The intracellular levels of oxidative stress were simulated based on the following formula:

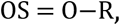

where O is the average production rate of all major oxidative molecules in a tissue, R is the antioxidant capacity activated in the same tissue, and OS represents the level of oxidative stress. We performed analyses for the three gene sets O, R, and S, and determined that their comprehensive expression levels reflecting the quantities O, R, and OS, respectively.

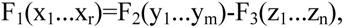

where r, m, and n are the number of genes in the three genomes, respectively, and equations F1, F2 and F3 were used to integrate the expression levels of selected genes for oxidative stress, oxidative molecule production, and antioxidant molecules, respectively. We referred to Xu Ying’s laboratory to construct polynomials to parameterize and validate the model. Finally, we applied a mature polynomial model to this data for the quantification of oxidative stress.

### Analysis of the metabolic processes of immune cells

We used the pathway activity algorithm. The pathway activity algorithm is as follows:

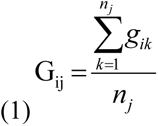

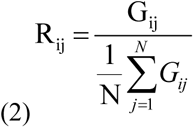

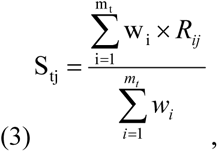

where Gij in equation (1) represents the average expression value of the i-th gene in the j-th cell type, nj represents the number of samples in the j-th cell type, and gik represents the expression value of the i-th gene in the k-th sample. In equation (2), Rij represents the relative expression value of the i-th gene in the j-th cell type. If Rij > 1, the average expression value of the gene in this cell is greater than the average expression value of the gene in all cells. N represents the number of cell types. In equation (3), Stj represents the score value of the j-th cell type in the t-th pathway, mt represents the number of metabolic genes in the t-th pathway, and wi represents the weight of the i-th gene.

## Acknowledgments

This work was supported by China Ministry of Science and Technology key research and development plan project [#2022YFF1203204]; The 2020 Biosafety Research Special Plan of the Logistics Support Department of the Military Commission[#923070201202]; Jilin Province Pathogen and Infection Informatics International Joint Research Center[#20210504004GH]; Young and middle-aged scientific and technological innovation leading talents and teams of Jilin Provincial Science and Technology Department[#20200301001RQ].

## Contributions

Design of research: Xizi Luo and Nan Zhang, Analysed data: Yuntao Liu, Beibei Du, Xuan Wang, Tianxu Zhao and Bingqiang Liu, Wrote the manuscript: Shishun Zhao, Jiazhang Qiu and Guoqing Wang.

## supplementary table

**Table 1:**
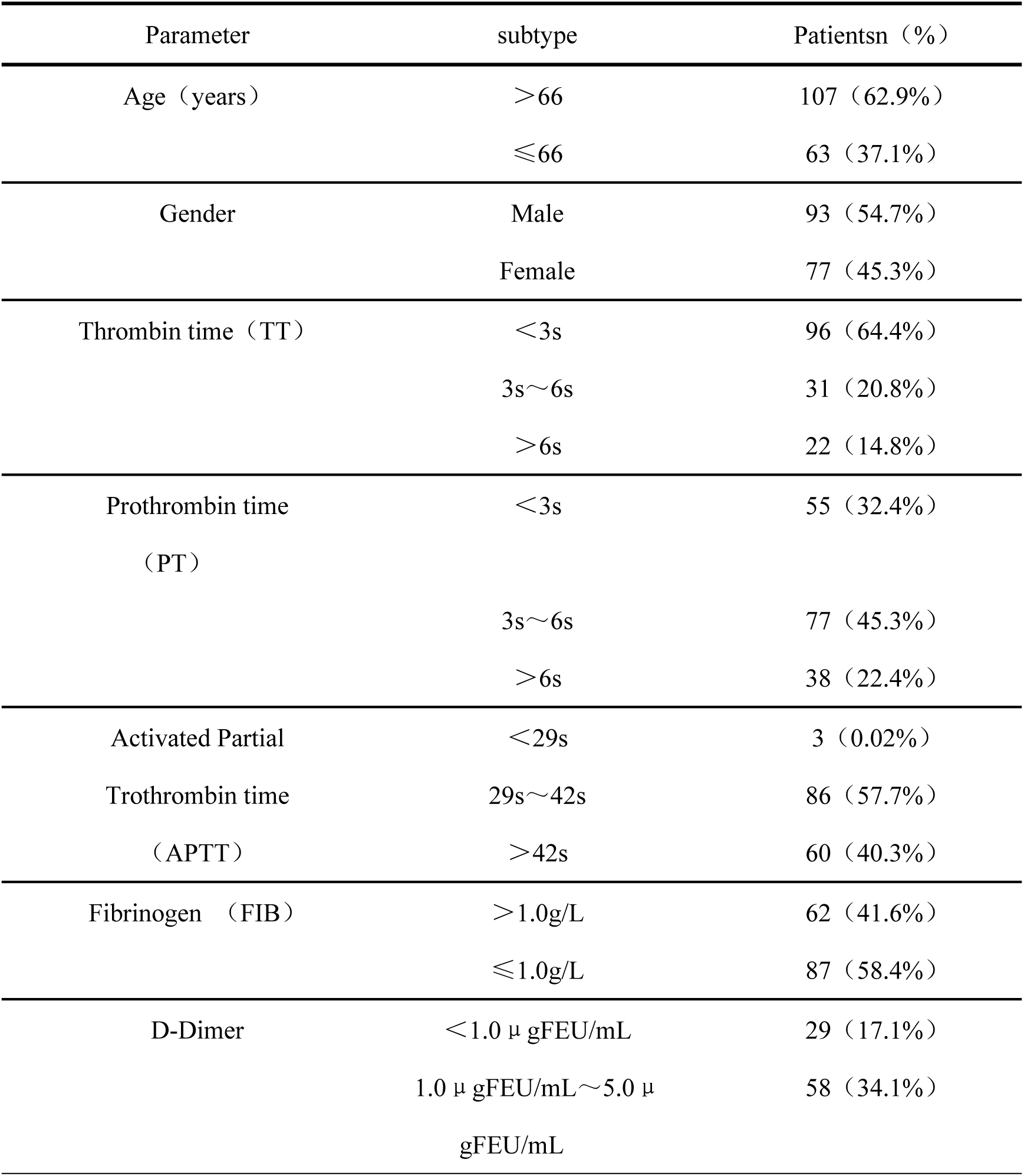
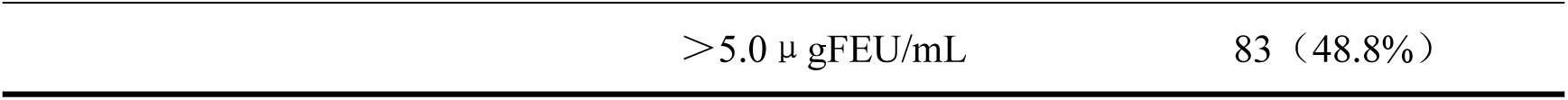
Analysis of coagulation factors in 170 critically ill patients with COVID-19

## Supplementary Figure

**Supplementary Figure 1.**
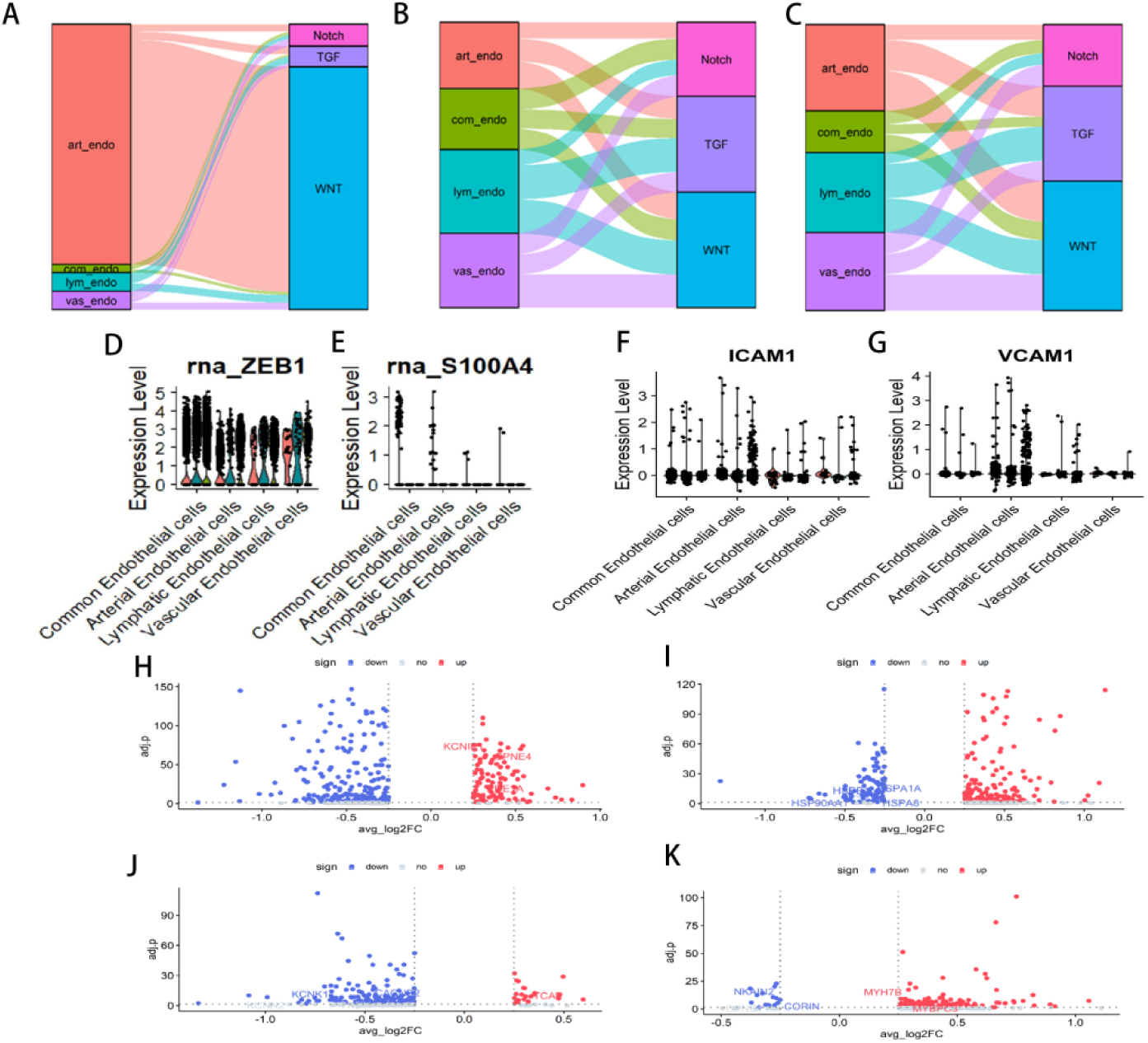
(A-C) Correlation analysis between endothelial cell subsets and Wnt, TGF, and Notch pathways during HC, CAC hypercoagulation, and CAC fibrinolysis (D-G) Expression of ZEB1, S100A4, ICAM1, and VCAM1 in endothelial cell subsets during HC, CAC hypercoagulation, and CAC fibrinolysis (H) Differential gene volcano map of ACTB^+^Cardiomyocytes in the hypercoagulable phase of CAC compared with HC (I) Differential gene volcano map of FHL2^+^Cardiomyocytes in the hypercoagulable phase of CAC compared with HC (J) Differential gene volcano map of ACTB^+^Cardiomyocytes in the CAC hypercoagulable phase compared with the CAC fibrinolytic phase (K) Differential gene volcano map of FHL2^+^Cardiomyocytes in the CAC hypercoagulable phase compared with the CAC fibrinolytic phase

**Supplementary Figure 2.**
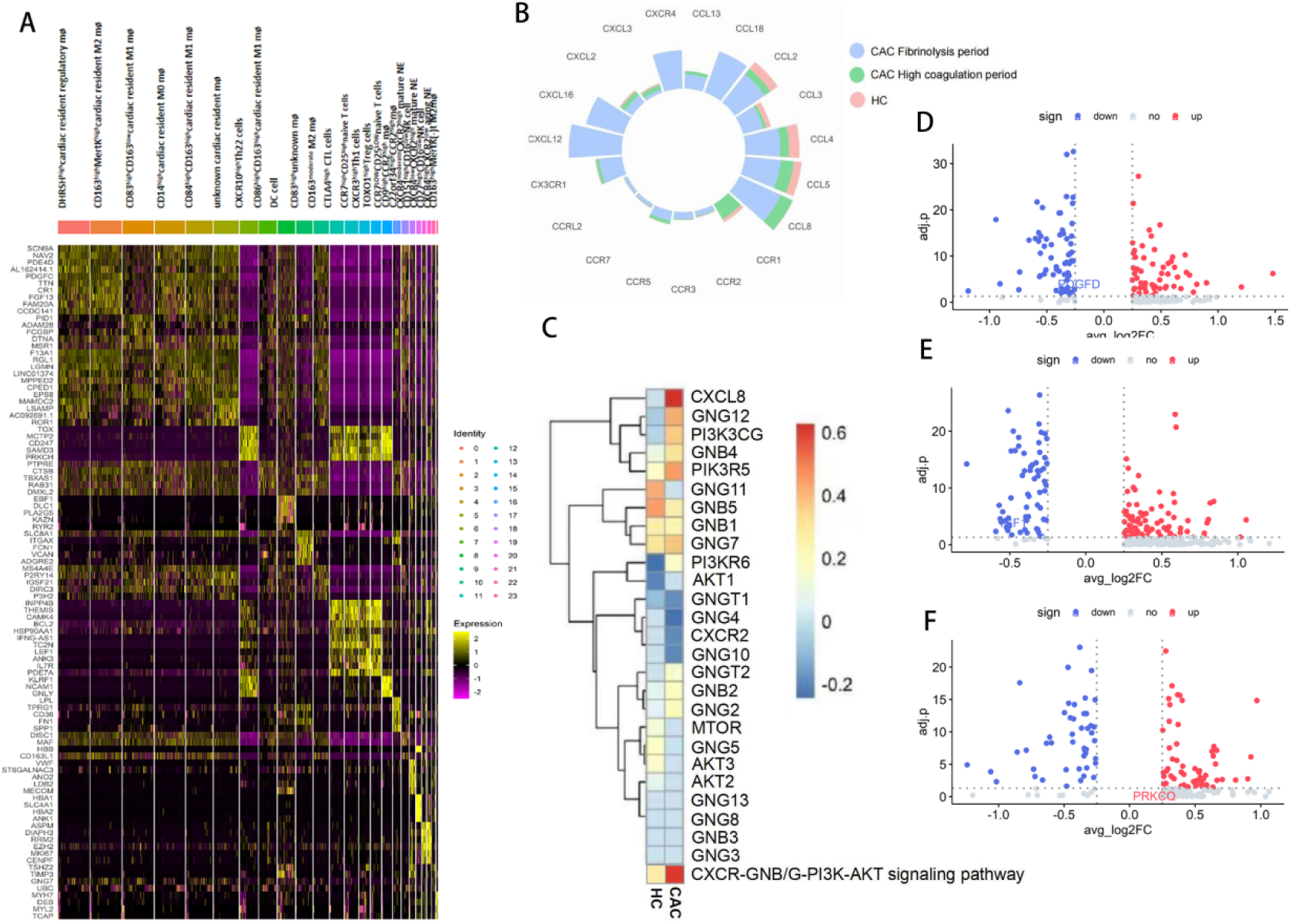
(A) Heatmap of selected marker genes of immune cell subsets within cell lineages (B) In HC, CAC hypercoagulable phase, and CAC fibrinolytic phase, the expression of selected chemokines (C) Correlation analysis between macrophages and the CXCR-GNB/G-PI3K-AKT signaling pathway in HC and CAC (D) CXCR10^high^Th22 cell differential gene volcano map in the CAC hypercoagulation period compared with the CAC fibrinolysis period (E) CD163^high^MERK^high^M2 macrophage differential gene volcano map in the CAC hypercoagulation period compared with the CAC fibrinolysis period (F) Native T cell differential gene volcano map in the CAC hypercoagulation period compared with the CAC fibrinolysis period

**Supplementary Figure 3.**
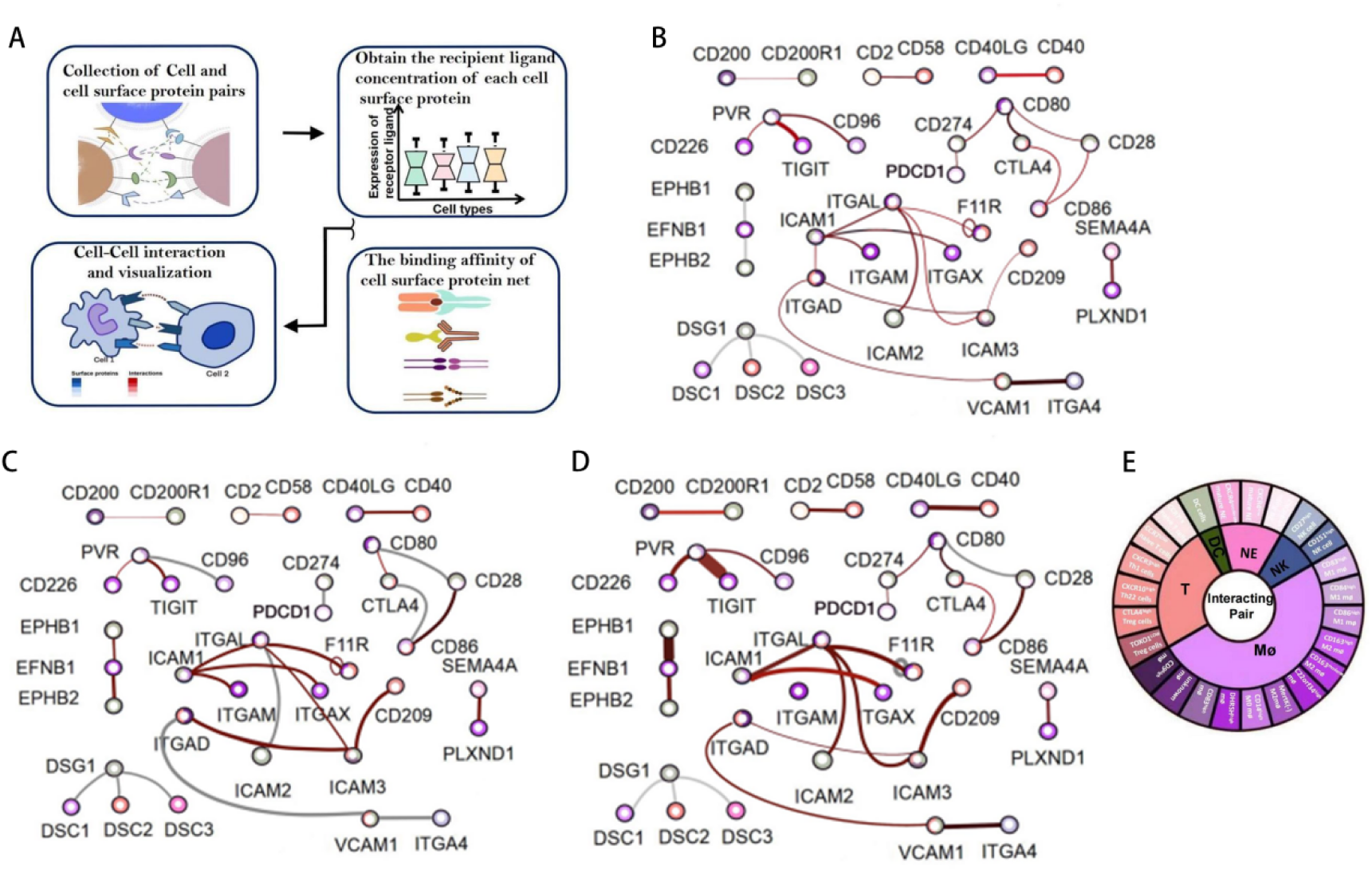
(A) Flow chart of surface protein network establishment (B-D) HC, CAC hypercoagulable phase, CAC fibrinolytic phase of the distribution of binding force of selected surface proteins (E) Schematic diagram of the color represented by immune cells in surface protein binding

**Supplementary Figure 4.**
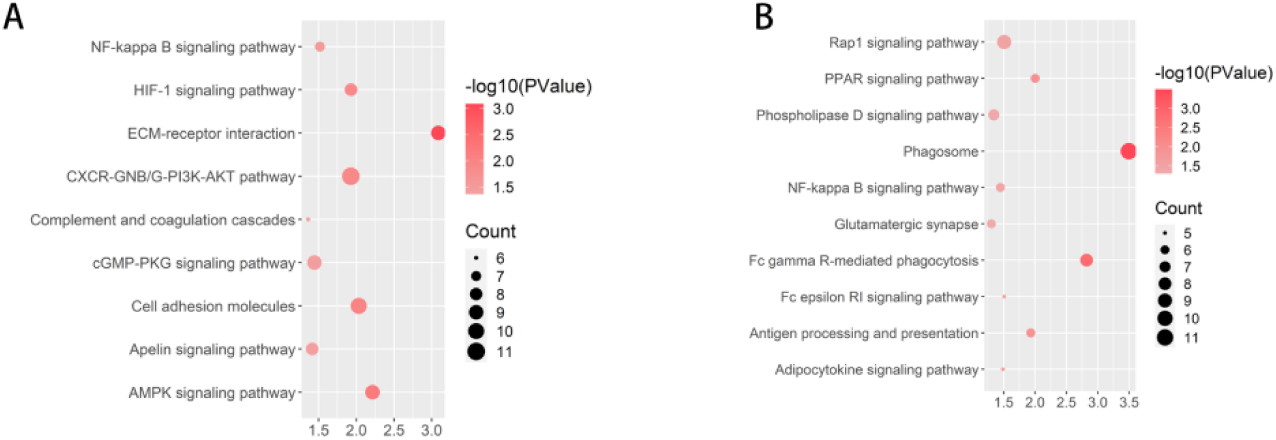
(A) Differential pathway enrichment in CD9^high^CCR2^high^monocyte-derived mø cells during the hypercoagulable phase of CAC compared with the fibrinolytic phase of CAC (B) Differential pathway enrichment in C22orf34^high^CCR2^high^monocyte-derived mø

**Supplementary Figure 5.**
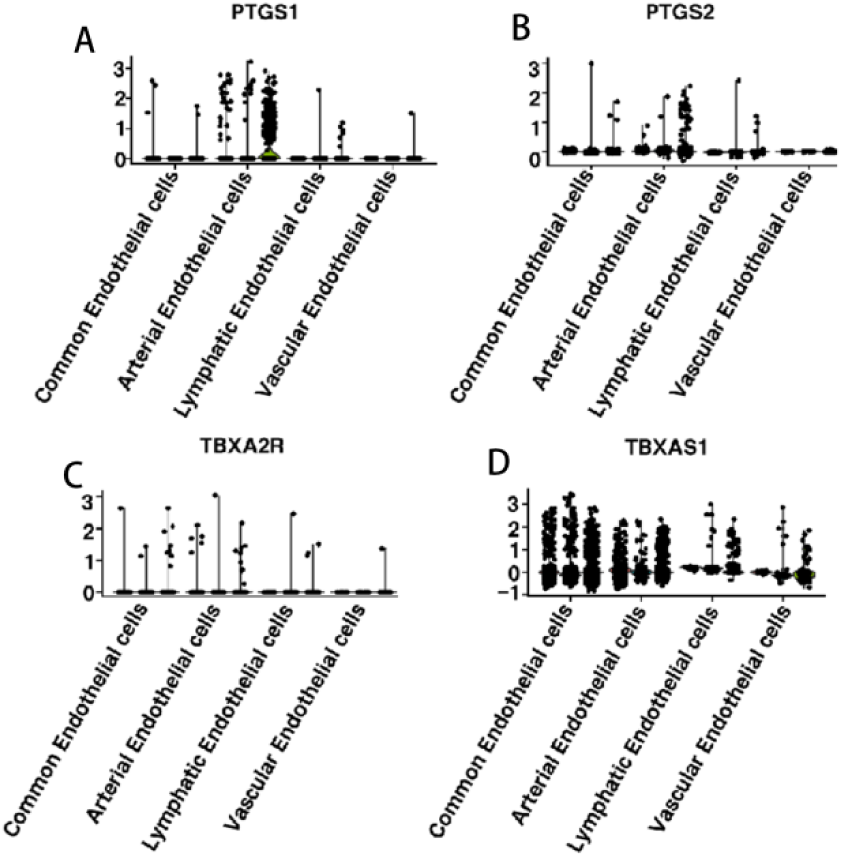
(A-D) The expression of PTGS1, PTGS2, TBXA2R, and TBXAS1 genes in endothelial cell subpopulations during HC and the CAC hypercoagulable and CAC fibrinolytic phases

